# A Simple and Reproducible In-Vivo Rabbit Phonation Model for Glottic Insufficiency

**DOI:** 10.1101/2022.02.14.480391

**Authors:** William M. Swift, Ian T. Churnin, Osama A. Hamdi, Andrew M. Strumpf, Heather A. Koehn, Patrick S. Cottler, James J. Daniero

**Author notes:** **Please send correspondence, reprint requests, and proofs to:** James J Daniero, MD, MS, Department of Otolaryngology – Head and Neck Surgery, University of Virginia Health System, P.O. Box 800713, Charlottesville, Virginia 22908-0713, 1-434-924-2040 (voice), 1-434-982-3965 (fax). **Level of Evidence:** N/A.

## Abstract

**Introduction:** Glottic insufficiency can result from neurologic injury, surgery, radiation, and the aging larynx. Treatment includes voice therapy, vocal fold injection augmentation, surgical medialization, or laryngeal reinnervation procedures. The objective of this study is to describe an in-vivo rabbit phonation model for glottic insufficiency that is simple and reproducible by means of unilateral cricothyroid muscle stimulation and high-speed video recordings of evoked phonation.

**Methods:** A non-randomized controlled trial utilizing seven New Zealand white rabbits was performed via a single operation including evoked phonation with bilateral and unilateral cricothyroid muscle stimulation conditions. The effect of stimulation method on glottic cycle, pitch and loudness was compared. Endoscopic recordings using 5,000 frames-per-second image capture technology and audiologic recordings were obtained for all phonation conditions. Primary outcome measures included means of maximum glottal area (MGA)/length pixel ratio, right and left amplitude/length pixel ratios, calculated cycle frequency, auditory recorded frequency, and maximum auditory intensity. Measurements were obtained via pixel counts using imageJ. Paired t-tests compared the average values obtained over five consecutive glottic cycles for each unilateral and bilateral stimulation during evoked phonation.

**Results:** Mean (median, IQR) MGA/length was significantly greater with unilateral, 20.30 (19.13, 10.97), vs bilateral, 9.62 (8.33, 2.58), stimulation (p=0.043). Mean frequency (median, IQR), 683.46 Hz (658.5, 197.1) vs 479.92 Hz (458.1, 112); (p=0.027) and mean maximum intensity, 83.5 dB (83.5, 1) vs 76.3 dB (74.5, 4); (p=0.013) were significantly reduced from bilateral to unilateral stimulation. There was no significant difference of mean right amplitude/length between bilateral and unilateral.

**Conclusion:** The described model demonstrates a simple and reproducible means of producing glottic insufficiency and represents a pathway for better understanding the biomechanics and pathophysiology of glottic insufficiency and offers the potential to compare treatment modalities through in-vivo study.

## INTRODUCTION

Glottic insufficiency is the incomplete approximation of the vocal folds during phonation and swallowing resulting in weak voice, difficulty communicating and aspiration. This condition often results from vocal fold paresis, paralysis, presbylarynges, or tissue loss due to surgical resection, trauma, and radiation. Treatment is varied depending on the etiology of insufficiency, but often involves voice therapy combined with surgery, such as injection augmentation, medialization laryngoplasty, or laryngeal reinnervation. Ultimately, there remains a significant role for the development of a representative model by which to expand and optimize research into the biomechanics, pathophysiology and treatments of glottic insufficiency.

Rabbit laryngeal experimental models possess multiple benefits including their similarity to human vocal fold histology and structure.^4, 5^ In addition, the rabbit larynx is large enough to easily facilitate surgical procedures and allow for the use of standard pediatric sized surgical instruments. The adult New Zealand white rabbit airway approximates the size of a human neonate. Furthermore, a rabbit in-vivo model of phonation proposed by Ge et al. demonstrated medialization of the vocal folds and phonations through bilateral stimulation of the cricothyroid muscles.^6^

Rabbit in-vivo phonation models have since been used for a variety of studies including the evaluation of physiologic effects of phonation on inflammatory markers^7–9^ and microflap wound healing.^10, 11^ Yet to date, the only small animal model simulating unilateral vocal fold paralysis (sUVFP) has been in cases where the recurrent laryngeal nerve (RLN) was sacrificed as a separate procedure.^12^ Utilizing RLN sacrifice as a survival surgery requires detailed microsurgery and is limited by the months of time required for subsequent vocalis muscle atrophy and is complicated by variable patterns or reinnervation limiting the reproducibility of this model.

We propose a simple modification of the existing Ge et al. protocol, creating a unique model for in-vivo simulation of glottal insufficiency due to unilateral vocal fold immobility. This model would allow for detailed analysis of the biomechanics of glottic insufficiency and vocal fold medialization/augmentation procedures in a single surgical procedure. The purpose of this study is to establish an in-vivo rabbit model for glottal insufficiency due to unilateral vocal fold immobility that is simple, reproducible via unilateral stimulation of the cricothyroid muscle.

## MATERIALS AND METHODS

In accordance with the University of Virginia Animal Care and Use Committee approved protocol (#4194), seven female New Zealand white rabbits (ranging 2.4-3.7kg) were used for this study. Each rabbit underwent evoked phonation with bilateral and unilateral right-sided cricothyroid muscle stimulation. In this manner, each rabbit served as its own control.

### Details of procedure

The rabbits were anesthetized with a combination of ketamine 50 mg/kg and xylazine 5 mg/kg given intramuscularly, with maintenance of sedation achieved via inhaled isoflurane (2.0 - 2.5%) and intravenous midazolam as needed. The larynx and trachea were exposed by a vertical midline neck incision extending from the hyoid bone to the sternal notch, and the larynx was suspended with a custom-modified pediatric Parsons infant laryngoscope with an aperture created on the bottom surface to allow for laryngeal access around the long maxillary central incisors of a rabbit. Two tracheotomies were performed. A distal tracheostomy was made adjacent to the sternum for placement of a size 3.0 cuffed endotracheal tube in the thoracic trachea for positive pressure ventilation and delivery of the isoflurane inhalant anesthetic for general anesthesia. Heated (37 °C) and 100% humidified airflow was delivered through the superior cervical tracheostomy to the glottis via size 3.5 cuffed endotracheal tube to entrain a sustained vocal fold mucosal wave. The cricothyroid muscles were isolated bilaterally, and pairs of 0.008” coated stainless steel anode-cathode wire electrodes (A-M Systems, part #791400, Sequim, WA) were inserted into each cricothyroid muscle. A pulse stimulator (A-M Systems, Model 2100, Sequim, WA) was used to deliver square wave pulses to each cricothyroid muscle. Vocal fold approximation during phonation was stimulated using a 60-Hz, 80-mA bipolar stimulus bilaterally in series configuration with a train of 1-ms pulses, with stimulus duration of 3 seconds on followed by 3 seconds off. Unilateral stimulation producing glottic insufficiency was performed with a 40-mA stimulation to only the right cricothyroid muscle. An airflow of 10L/min was used to sustain vocal fold oscillation in both conditions. An image of the experimental procedure set up is shown in Figure 1.

**Figure 1:**
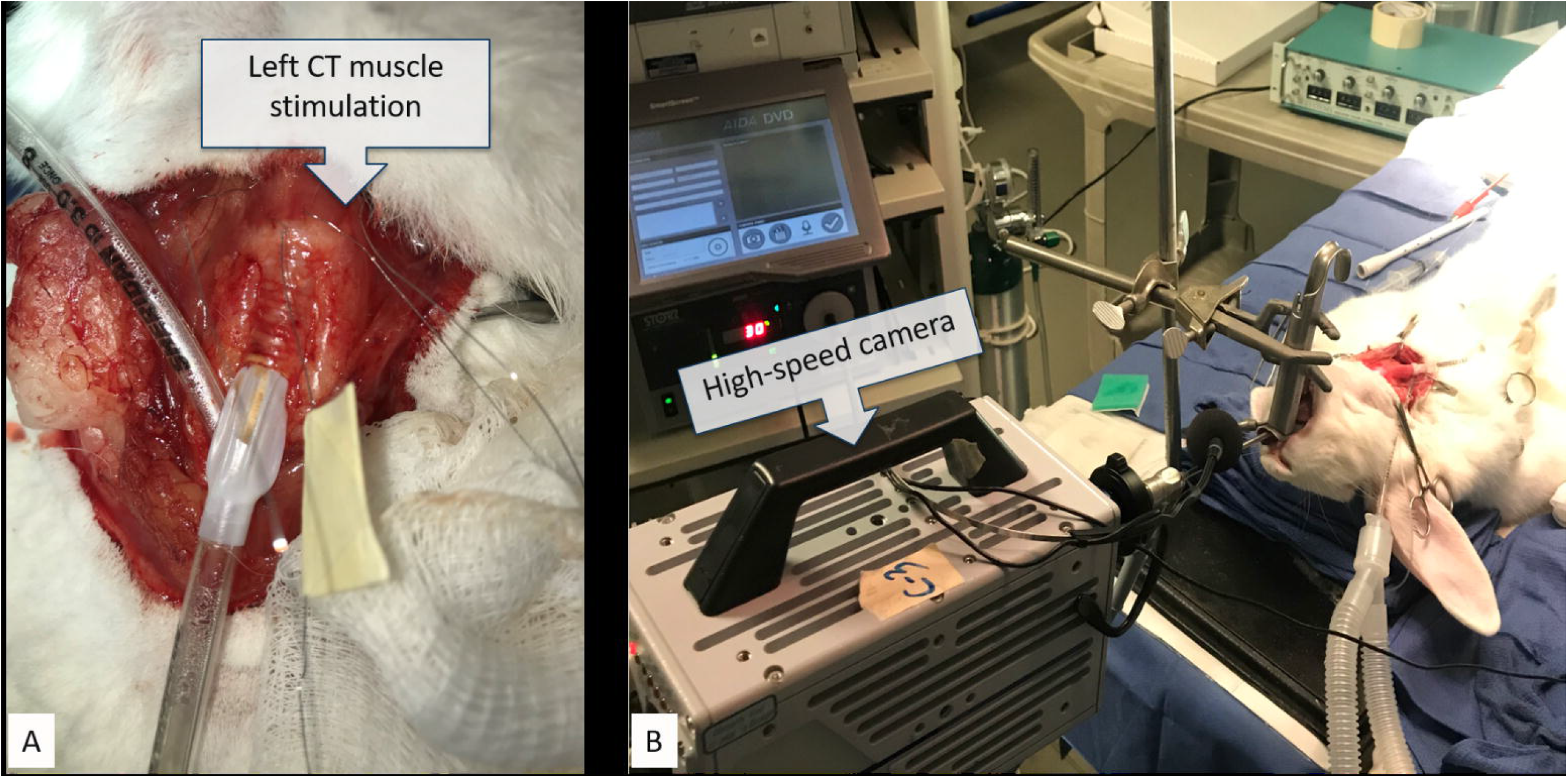
A representative image of the experimental procedure set up. In rabbits the cricothyroid muscle is responsible for vocal fold medialization. Stimulation of the right cricothyroid muscle alone simulated unilateral (left) vocal fold paralysis (figure 1A). A high speed camera was used to capture video of the glottis during phonation (figure 1B).

Image capture during phonation was recorded using a high-speed video camera (Fastec Imaging IL-5M, San Diego, CA) in combination with a 3.0-mm, 30° rigid endoscope (KARL STORZ Endoskope, El Segundo, CA). Video parameters included recording at 5000 frames per second in monochrome, shutter speed of 187 microseconds and a gain of 1.0 with spatial resolution of 256 x 512 pixels. 2X binning and 10 bit image settings were used for image processing using FasMotion version 2.5.3.

In addition to image capture, audio recording and processing during phonation was performed using Voice Analyst 3.8 via iOS 14.3 and iPhone12 Pro microphone. Acoustic parameters recorded included average pitch and intensity.

### Data Analysis

Analysis of the captured images was performed for phonation cycles during stable periodic mucosal waves in the mid-portion of the three second phonation that were representative of the entire phonation trial. To assess glottic competence key measurements were obtained from still images. Five successive cycles of vocal fold mucosal wave were selected during the mid-portion of sustained phonation. Maximal glottal area was measured during peak amplitude using frame-by-frame analysis. A representative image used for data analysis is displayed in Figure 2. The image with the maximal glottic opening for each of the five cycles was analyzed with ImageJ 1.53 (National Institutes of Health, Bethesda, MD). Measurements determined with ImageJ included, the pixel area of glottic opening corrected for vocal fold length, the amplitude defined as the distance from midline to vocal fold at the mid-membranous portion of the vocal fold on both the left and right sides via pixel count. Maximal glottic area as well as left and right amplitude were divided by the glottic length to standardize images and allow for comparison between phonation trials that could vary in endoscope positioning. Pitch was calculated mathematically from the video for measured cycles based on the known rate of 5000 images per second.

**Figure 2:**
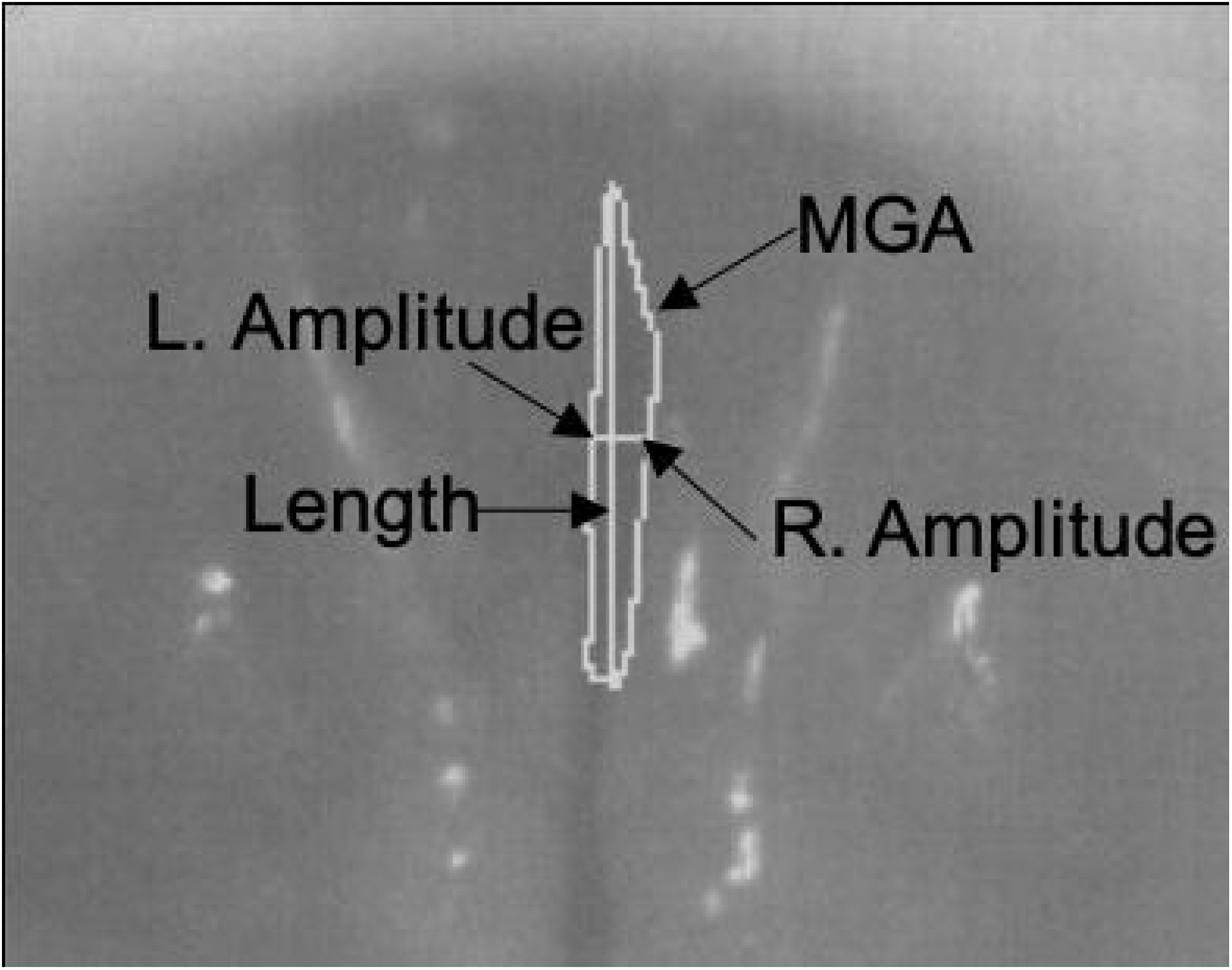
A representative image of the high-speed images obtained of the glottis along with key measurements used for data analysis.

Calculated pitch was compared to audio recorded maximum pitch. Peak phonation pitch was achieved during stable phonation eliminating the ramp and taper associate with onset and offset of phonation. The measurements for the five cycles during bilateral and unilateral stimulation were each averaged (mean) for each rabbit. All measurements were performed by a single rater that was trained by the senior author (J.J.D.) and blinded to the conditions.

### Statistical analysis

Variables included in the analysis were the MGA/length pixel ratio, amplitude (right and left) per length pixel ratio, recorded maximal pitch, calculated pitch, and maximal volume intensity. The five measured pixel ratio values for each variable were averaged (mean) for each rabbit, and then values for all the rabbits were averaged (mean) and compared for bilateral vs unilateral cricothyroid stimulation conditions. Shapiro-Wilk test was used to determine normality within the data sets. Paired t was used to test for statistical significance with a p-value cut off of p < 0.05 indicating significance.

## RESULTS

Seven rabbits underwent surgery using the glottic insufficiency phonation model. One rabbit suffered a laryngeal hematoma that immobilized the right vocal fold and prevented bilateral or unilateral phonation, leading to its exclusion from data analysis. The remaining six rabbits underwent the phonation procedure without complication.

The mean MGA/Length for bilateral stimulation of all rabbits was 9.62 compared to 20.30 (p = 0.043) during unilateral stimulation as depicted in Figure 3a. There was not a significant difference in comparing the amplitude of the right or left vocal cord under bilateral vs unilateral stimulation, 0.11 compared to 0.06 with bilateral stimulation(p=0.059). The results for the right and left sided amplitude are shown in Figures 3b and 3c, respectively.

**Figure 3A, B, C:**
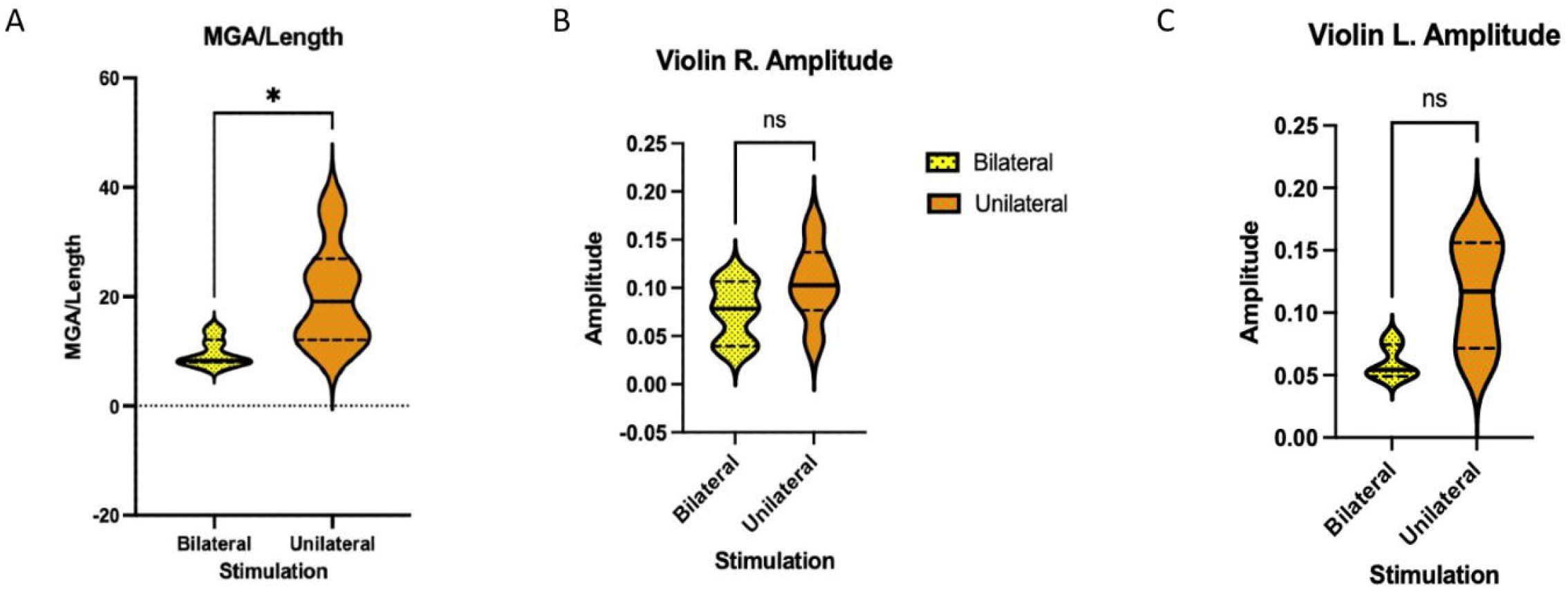
The mean maximum glottal area/length for bilateral stimulation of all rabbits compared with unilateral stimulation (figure 3A). The amplitude of the right vocal cord under bilateral and unilateral stimulation (figure 3B) compared with the left vocal cord (figure 3C).

Bilateral stimulation resulted in a higher pitch frequency when determined by audio recording and when calculated via high-speed image capture as shown in Figure 4. The mean pitch frequency using acoustic analysis was 638.67 Hz under bilateral stimulation compared to 410.17 Hz during unilateral stimulation (p=0.020). Similarly, when calculating frequency via image capture, bilateral stimulation resulted in a mean pitch of 683.46 Hz compared to 479.92 Hz under unilateral stimulation (p=0.027). Additionally, there was no significant difference in the values obtained using the recorded versus calculated methods of determining frequency for either bilateral (p=0.62) or unilateral recorded vs calculated (p=0.467).

**Figure 4:**
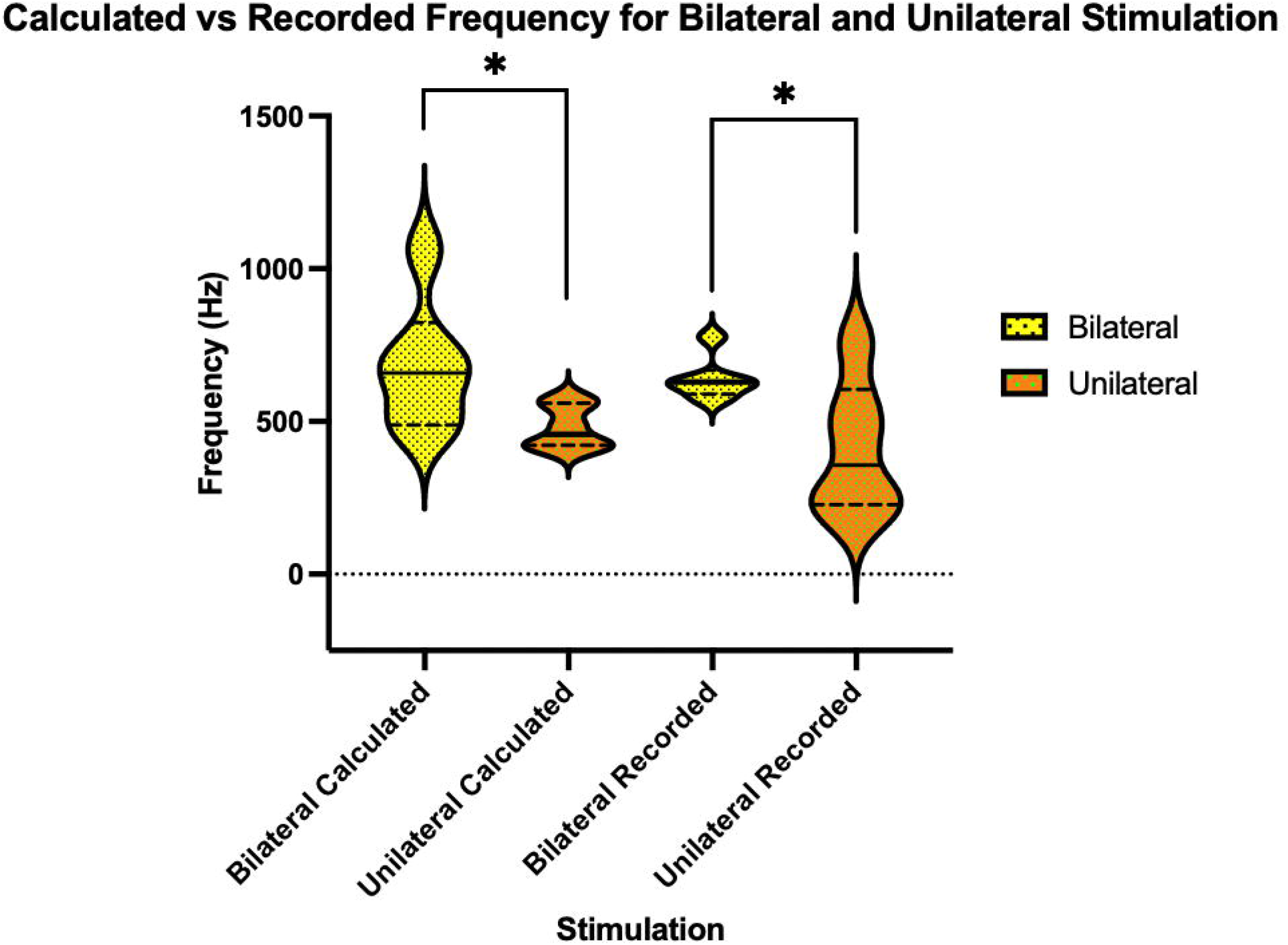
Frequency of vocal cords under bilateral and unilateral stimulation was calculated via image capture and compared with recorded frequencies.

In comparing the maximal intensity during vocal cord stimulation, bilateral stimulation resulted in a significantly higher intensity than with unilateral stimulation. The mean maximal intensity under bilateral stimulation was 83.5 dB compared to 76.3 dB during unilateral stimulation (p=0.013) as depicted in Figure 5. This change represents nearly a ten-fold decrease in sound intensity due to only unilateral stimulation. The results of evoked phonation analysis are summarized in table 1.

**Table 1:** Control (Bilateral) vs. Glottic insufficiency model (unilateral), glottic insufficiency, frequency, and intensity analysis

| Outcome | Cricothyroid Stimulation |  |  |  |  |  |  |  |  |
| --- | --- | --- | --- | --- | --- | --- | --- | --- | --- |
|  | Bilateral |  |  |  | Unilateral |  |  |  |  |
|  | Mean | Median | IQR | SD | Mean | Median | IQR | SD | P |
| **MGA/length** | 9.62 | 8.33 | 3.37 | 2.52 | 20.3 | 19.13 | 11.74 | 9.35 | **0.0428\*** |
| **Amplitude (R)** | 0.075 | 0.078 | 0.066 | 0.031 | 0.105 | 0.103 | 0.041 | 0.039 | 0.208 |
| **Amplitude (L)** | 0.060 | 0.054 | 0.023 | 0.013 | 0.114 | 0.117 | 0.076 | 0.043 | 0.0592 |
| **Calculated Pitch (Hz)** | 683.5 | 658.5 | 255.4 | 212.01 | 479.9 | 458.1 | 129.4 | 70.93 | **0.027\*** |
| **Recorded Max Pitch (Hz)** | 638.7 | 629 | 36 | 72.18 | 410.2 | 356.5 | 315 | 219.3 | **0.020\*** |
| **Maximal Volume (dB)** | 83.5 | 83.5 | 1 | 1.64 | 76.3 | 74.5 | 5 | 4.97 | **0.013\*** |

MGA: Maximum glottic area

**Figure 5:**
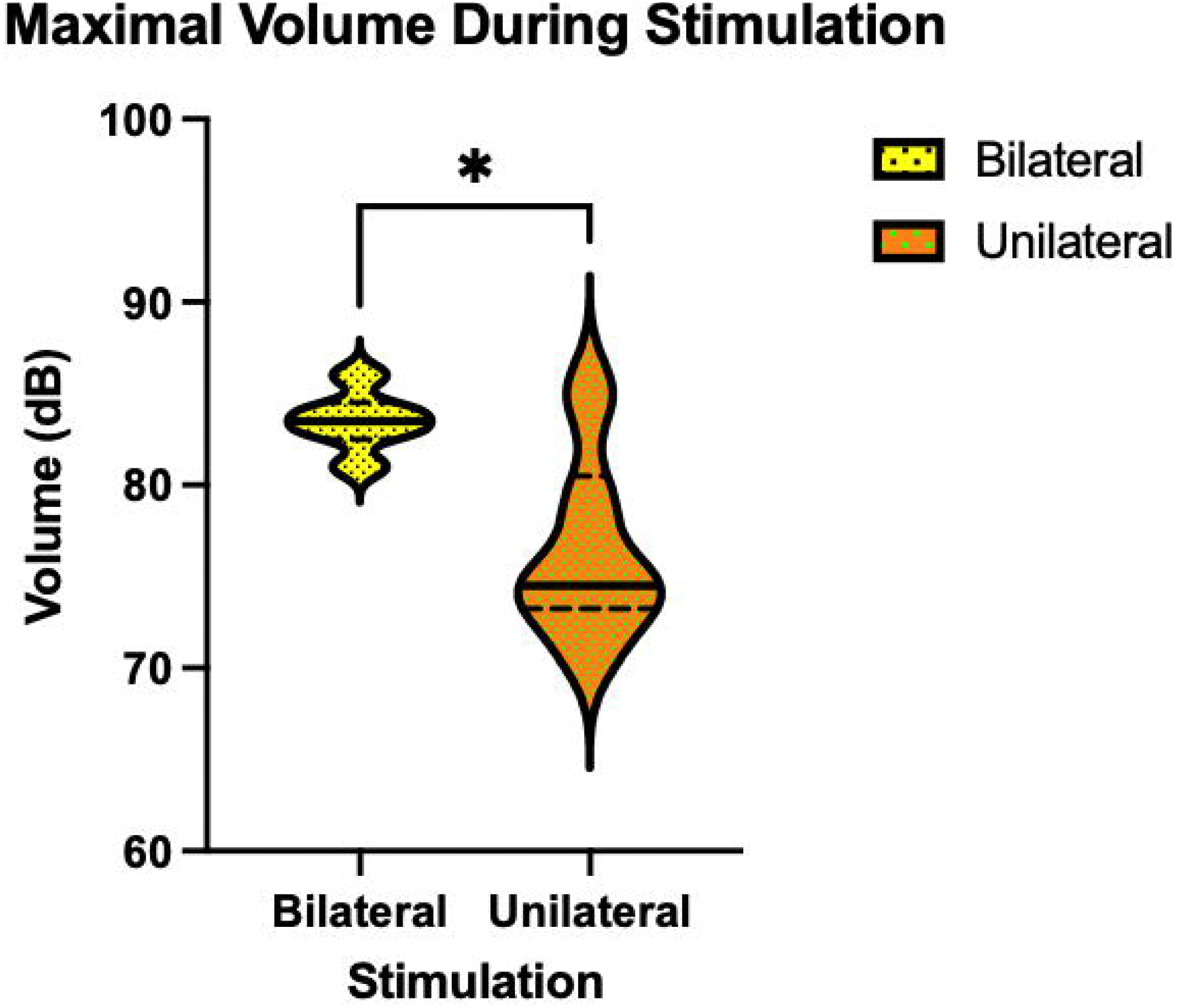
The maximal intensity during both bilateral and unilateral vocal cord stimulation.

## DISCUSSION

Glottic insufficiency is a pathologic process that can carry a substantial burden on health, social, and occupational quality of life due to weak voice production and liquid aspiration.^13, 14^ The described model may provide a simple alternative for sUVFP research with the potential for rapid iteration because there is no need for laryngeal denervation and it does not alter the laryngeal framework.

Previous phonation of rabbit larynges have been demonstrated by Ge et al. In their study, glottic closure was achieved via bilateral stimulation of the cricothyroid muscles at 3-4 mA and 50-60 Hz.^6^ This method has served as the basis for performing computational flow modeling,^15^ and studying vocal fold changes due to phonation.^8, 9^ More recently, Li et al. were able to model asymmetric vocal fold closure characteristic of UVFP via a unilateral cricothyroid suture approximation.^16^

We demonstrated the feasibility of a similar in-vivo rabbit phonation model utilizing a method of unilateral stimulation to reproduce glottic insufficiency. Our model uniquely relies on selective unilateral vocal fold stimulation which offers an alternative to the surgical modification performed by Li et al.^16^ that alters the laryngeal framework. Variability in suture placement and tension can alter results from rabbit to rabbit using the unilateral cricothyroid approximation model; however, similar laryngeal configurations can be achieved via unilateral electrical stimulation with precise stimulation settings to achieve reproducible results in producing glottic insufficiency.

Our in-vivo cricothyroid simulation model produced data representative of a glottic insufficiency. As expected, the analysis of MGA (standardized by vocal fold length) demonstrated a significantly larger glottic area in the unilateral compared to the bilateral cricothyroid stimulation phonations (20.30 compared to 9.62, p = 0.043). This is due to glottic insufficiency from the lateral resting position of the left vocal fold in the non-stimulated side during unilateral stimulation. This enlarged glottic gap during phonation is the key finding that demonstrates the effectiveness of our model. The analysis of left amplitude represented an assessment of the stimulated compared to the non-stimulated true vocal fold. Expected results were an increased left vocal fold amplitude due to reduced tone on the unstimulated side. The comparison of left amplitude between the two groups did not reveal a statistically significant difference, but with n=6, this parameter could be more subtle and require more rabbits to resolve the difference. The analysis of the right amplitude represented an assessment of the true vocal fold that was stimulated in both groups and served as a control. As anticipated, there was no significant difference in the right true vocal fold amplitude between the unilateral and bilateral stimulation groups.

With the known 5,000 frames per second image capture, the review of the sequential glottic cycle images allowed for the calculation of the duration of each glottic cycle, and therefore allowed for a calculated frequency (Hz) for each cycle. In order to assess validity of this ideal objective measurement of pitch, the calculated frequency was compared to recorded frequency for both unilateral (p = 0.467) and bilateral stimulation (p = 0.62). The results demonstrated no significant difference, supporting the use of calculated frequency for future studies as an objective means of determining pitch effects with the glottic insufficiency model. Finally, maximum volume intensity achieved with bilateral stimulation was significantly greater than with unilateral stimulation.

Ultimately, our proposed model effectively demonstrates that unilateral cricothyroid stimulation simulates glottic insufficiency due to unilateral vocal fold immobility. In addition, and perhaps most critically, this model would represent a single surgery means of assessing glottic insufficiency that would obviate the need for more complex staged recurrent laryngeal nerve transection methods that have been previously described and used in research for treatments for glottic insufficiency.^12, 17, 18^ Furthermore, this reproducible technique avoids the variability of reinnervation patterns introduced with use of the recurrent laryngeal nerve section. In addition to the reduced burden of prolonged animal housing and a second surgical phonation surgery needed in recurrent laryngeal nerve sectioning methods, our method will limit the number of animals sacrificed as each rabbit can serve as both a case and control subject. The 3 R’s of animal research (reduction, refinement, and replacement) are upheld with this streamlined protocol for glottic insufficiency. This model improves on the “refinement” of the previous procedure by limiting the animal to a single surgical procedure without paralysis, and minimizes the discomfort to the animal. This phonation model, utilizing cricothyroid stimulation as a means of objectively assessing glottic function through measured outcomes such as maximum glottic area, amplitude, calculated frequency, and vocal intensity could be translated into research evaluating other etiologies of glottic insufficiency.

The research and clinical implications of this alternative model are valuable as it represents a pathway for further reproducible studies to better understand glottic insufficiency. Our lab aims to further characterize study this model using computation flow dynamics. The proposed model would be an ideal method to conduct head-to-head comparison of different phonation conditions in the setting of glottic insufficiency.

**CONCLUSION**

The proposed in-vivo rabbit phonation model utilizing cricothyroid stimulation represents a single surgical and reproducible means to simulate glottic insufficiency with measurable objective outcomes including maximal glottic area, phonation frequency and sound intensity. With the inherent limitations and nature of animal-based research, this model represents a promising pathway for better understanding glottic insufficiency and expediting the development of novel treatment modalities. This model will hopefully allow for detailed analysis of the biomechanics of glottic insufficiency and head-to-head assessment of vocal fold procedures in a single standardized surgical procedure.

